# Transdermal electrical neuromodulation of the trigeminal sensory nuclear complex improves sleep quality and mood

**DOI:** 10.1101/043901

**Authors:** Alyssa M. Boasso, Hailey Mortimore, Rhonda Silva, Linh Aven, William J. Tyler

## Abstract

Achieving optimal human performance that involves cognitive or physical work requires quality sleep and a positive mental attitude. The ascending reticular activating system (RAS) represents a powerful set of endogenous neuromodulatory circuits that gate and tune global brain responses to internal and external cues, thereby regulating consciousness, alertness, and attention. The activity of two major RAS nuclei, the locus coeruleus (LC) and pedunculopontine nucleus (PPN), can be altered by trigeminal nerve modulation. Monosynaptic afferent inputs from the sensory components of trigeminal nerve branches project to the trigeminal sensory nuclear complex (TSNC), which has direct and polysynaptic connections to the LC and PPN. We previously found high-frequency (7 – 11 kHz) transdermal electrical neuromodulation (TEN) of the trigeminal nerve rapidly induces physiological relaxation, dampens sympathetic nervous system responses to acute stress, and suppresses levels of noradrenergic biomarkers. Given the established roles of LC and PPN neuronal activity in sleep regulation, psychophysiological arousal, and stress, we conducted three studies designed to test hypotheses that modulation of the TSNC can improve sleep quality and mood in healthy individuals (n = 99). Across a total of 1,386 days monitored, we observed TEN modulation of trigeminal and cervical nerves prior to sleep onset produced significant improvements in sleep quality and affective states, quantified using clinically validated surveys, overnight actigraph and heart rate recordings, and biochemical analyses compared to baseline or sham controls. Moreover, we observed some frequency dependence in that TEN delivered at lower frequencies (TEN_LF_; 0.50 – 0.75 kHz) was significantly more effective at improving sleep quality and reducing anxiety than higher frequency TEN waveforms. Collectively our results indicate that transdermal electrical neuromodulation of trigeminal and cervical nerve branches can influence TSNC activity in a manner that significantly improves sleep quality and significantly reduces stress. We conclude that biasing RAS network activity to optimize sleep efficiency and enhance mood by electrically modulating TSNC activity through its afferent inputs holds tremendous potential for optimizing mental health and human performance.

## INTRODUCTION

The ascending reticular activating system (RAS) is a collection of nuclei and circuits that sort, filter, integrate, and transmit incoming sensory information from the brain stem to the cortex to regulate sleep/wake cycles, arousal/alertness, attention, and sensorimotor behaviors [1-8]. The endogenous neuromodulatory actions of the RAS on consciousness and attention are orchestrated by at least three distinct sets of brain stem nuclei that include cholinergic neurons of the pedunculopontine nucleus (PPN), noradrenergic neurons of the locus coeruleus (LC), and serotonergic neurons of raphe nuclei [1]. Through a cytoarchitectural meshwork of interconnected brain stem nuclei in the pons and midbrain, sensory inputs first act upon the brain to engage ascending RAS networks, which generate global arousal (“waking”), alerting, and orienting cues as parsed sensory information projects through thalamic pathways onto the cortex for additional processing and integration. More specifically the local and distal synaptic circuits formed by axons of neuromodulatory RAS networks (including neurons of the LC, PPN, RN) gate information flow from the sensory environment to the cortex and, in an activity-dependent manner, are capable of rapidly triggering neurobehavioral transitions across different states of behavioral awareness and consciousness [7, 9-11]. Depending on their firing rates for example, neurons of the PPN can differentially mediate REM sleep states [1, 12] and neurons of the LC can trigger sleep/wake transitions [13, 14]. Disrupted activity of ascending RAS networks underlies several neuropsychiatric conditions and disorders, such as insomnia, anxiety, depression, post-traumatic stress disorder (PTSD), and attention deficit hyperactivity disorder (ADHD) [1, 15-17]. Therefore, a neural interface capable of dynamically and electrically modulating RAS networks should be able to provide a chemical-free approach to restoring poor daily function attributable to sleep loss or attention and mood disorders. Such an interface would also support new approaches capable of optimizing normal human performance.

The trigeminal nerve or the fifth cranial nerve (cranial nerve V) bilaterally innervates the anterior half of the head and face including around the top of the scalp, the forehead, around eye orbits, nasal region, lips, jaw, and the oral cavity. Three main branches of the trigeminal nerve (ophthalmic, maxillary, mandibular branches) and their thousands of sub-branches transmit sensory information (chemical, thermal, mechanical, pain, and proprioceptive) via monosynaptic connections to the trigeminal sensory nuclear complex (TSNC). The TSNC itself is an elongated structure with several functional divisions (for example, the primary sensory nucleus and the spinal nucleus) spanning from the cervical spinal cord to the midbrain. The TSNC has been functionally mapped using multimodal trigeminal stimulation combined with fMRI and DTI in humans [18, 19]. The TSNC also receives some non-trigeminal sensory information from the neck via cervical afferents [20]. In turn the TSNC projects incoming sensory information through ascending pathways to multiple brain regions that regulate arousal and coordinate neurobehavioral engagement with the environment, such as the thalamus [21, 22], the superior colliculus [21, 23, 24], the cerebellum [23, 25], and the inferior olive [23, 26]. Several other electrophysiological and neuroanatomical studies have provided definitive evidence of functional synaptic connectivity between trigeminal afferents and the PPN and LC [27-30].

Besides its robust functional connectivity to ascending RAS networks, a major advantage of the TSNC as a neuromodulation target is that its primary monosynaptic inputs can be noninvasively accessed and coupled to using safe and comfortable transdermal neurostimulation approaches. Transcutaneous trigeminal nerve stimulation (TNS) has been shown to be effective for treating neuropsychiatric conditions like depression [31], PTSD [32], generalized anxiety disorder (GAD)[33], ADHD [34], as well as neurological disorders like epilepsy [35, 36] and headache [37, 38]. Interestingly, acute TNS at a frequency of 120 Hz has been demonstrated to induce sleepiness and sedative like effects in healthy adults while 2.5 Hz stimulation frequency did not [39]. The observations made by Piquet and colleagues (2011) suggest that TNS may provide some benefit for some sleep disturbances and insomnia.

We recently described an approach to transdermal electrical neuromodulation of trigeminal and cervical nerve afferents that suppressed physiological and biochemical signatures of sympathetic tone and noradrenergic activity [40]. We showed that pulsed (7 - 11 kHz) TEN of trigeminal and cervical afferents significantly suppressed basal sympathetic tone compared to sham as indicated by functional infrared thermography of facial temperatures, significantly lowered levels of tension and anxiety on the Profile of Mood States scale compared to sham, and in response to acute stress induction TEN significantly suppressed changes in heart rate variability, galvanic skin conductance, and salivary α-amylase levels compared to sham [40]. Based on these findings, we hypothesized that repeated daily dampening of psychophysiological and biochemical arousal might improve mood if used nightly to increase the quality, duration or efficiency of sleep. Supportive of this hypothesis there are numerous lines of evidence that suggest insomnia is a “waking” disorder (hyper-arousal) of RAS networks rather than a sleep disorder *per se* [1, 41]. Viewed as such, we more specifically hypothesized that decreasing neurobehavioral and psychophysiological arousal by perturbing trigeminal LC/RAS networks prior to sleep onset for repetitive nights can enhance the restorative qualities of sleep on mood and mental health. Below we describe the results from our findings that demonstrate TEN of trigeminal afferents significantly improves sleep quality and mood on weeklong time scales. We speculate noninvasive modulation of the TSNC will provide a valuable platform approach to restoring and optimizing brain function and mental health.

## RESULTS

### Repeated nightly TEN significantly improves waking mood and reduces weekly stress compared to baseline

In Experiment 1, we examined the impact of before bed TEN on mood and mental health. Volunteers (n = 38) completed a week of baseline assessments followed by a week of nightly TEN treatments (20 min) self-administered within 30 min of going to bed (**Figure 1A**). Each morning, participants reported their mood using the Positive and Negative Affectivity Scale (PANAS) and by rating their degree of waking drowsiness and refreshment (See Methods). Mental health was captured by weekly responses to the Depression, Anxiety and Stress Scale (DASS). The PANAS and DASS are validated and widely used scales that capture fluctuations in mood and indicators of mental health, respectively.

**FIGURE 1.**
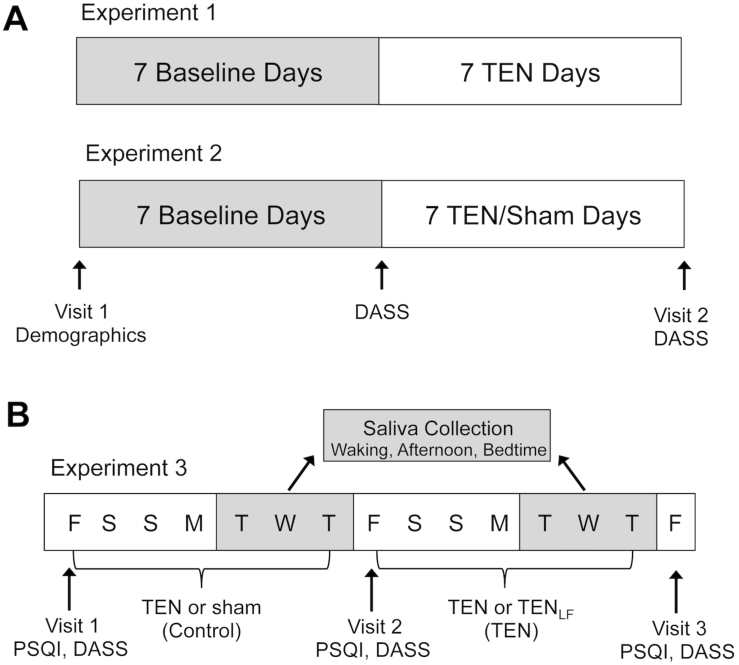
Experimental designs implemented for testing the effects of nightly repeated, self-administered transdermal electrical neuromodulation of trigeminal and cervical nerve sensory afferents on sleep and mood. A, The illustrations show that in Experiments 1 and 2 volunteers had one week or baseline assessment then self-administered transdermal electrical neuromodulation of trigeminal and cervical nerve afferents or an indistinguishable active sham waveform nightly before bed during the second week of testing. Several metrics were used to evaluate morning mood, weekly mental health indicators, and sleep quality (see Methods). **B**, In Experiment 3 we implemented a double-blinded crossover design where subjects self-administered an active sham treatment or real TEN (3 – 11 kHz, 5 – 7 mA average current amplitude) treatment nightly before bed for one week and then either real TEN or low-frequency TEN (TEN_LF_; 0.50 – 0.75 kHz, < 5 mA average current amplitude). Again an array of metrics were used to determine the impact of sham treatment, TEN-treatment, and TEN_LF_-treatment on sleep quality and mood.

Relative to baseline, TEN treatment improved positive affect by 9% (t(35) = 3.427, p = 0.002) and reduced negative affect by 10% (t(35) = −3.147, p = 0.003) according to the PANAS (Table 1). Compared to baseline, indicators of mental health recorded by the DASS (Table 1) revealed that TEN significantly reduced stress (t(30) = −3.982, p < 0.001) and anxiety (t(29) = −2.177, p = 0.038) by 41.3% and 30% respectively, whereas depression scores remained unchanged (p = 0.469; **Figure 2A**). Further, during the TEN treatment period, participants felt 27% less drowsy (t(36) = −4.859, p < 0.001) and 17.92% more refreshed (t(36) = 4.908, p < 0.001) upon waking compared to the baseline period (**Figure 2B** and Table 1). These data indicate that nightly TEN treatments self-administered prior to bed improve morning mood, awakening affect and arousal, and reduces stress and anxiety on weeklong time scales. The enhanced morning affectivity and arousal combined with the mechanisms of action posited by Tyler et al (2015) suggest TEN may enhance sleep quality when used prior to bedtime.

**Table 1.**
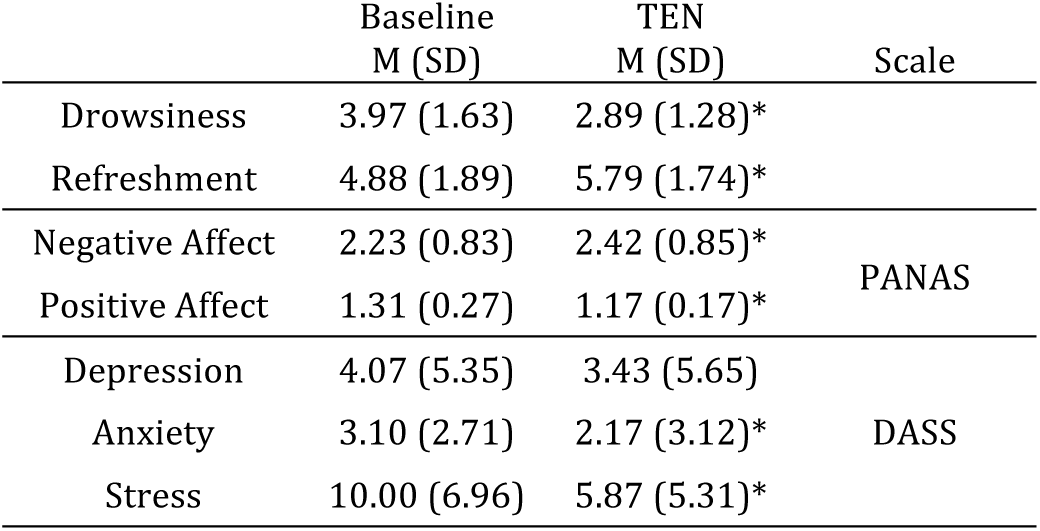
Summary of results from Experiment 1 illustrating the effects of nightly TEN on morning arousal, affect, and mental health.

**FIGURE 2.**
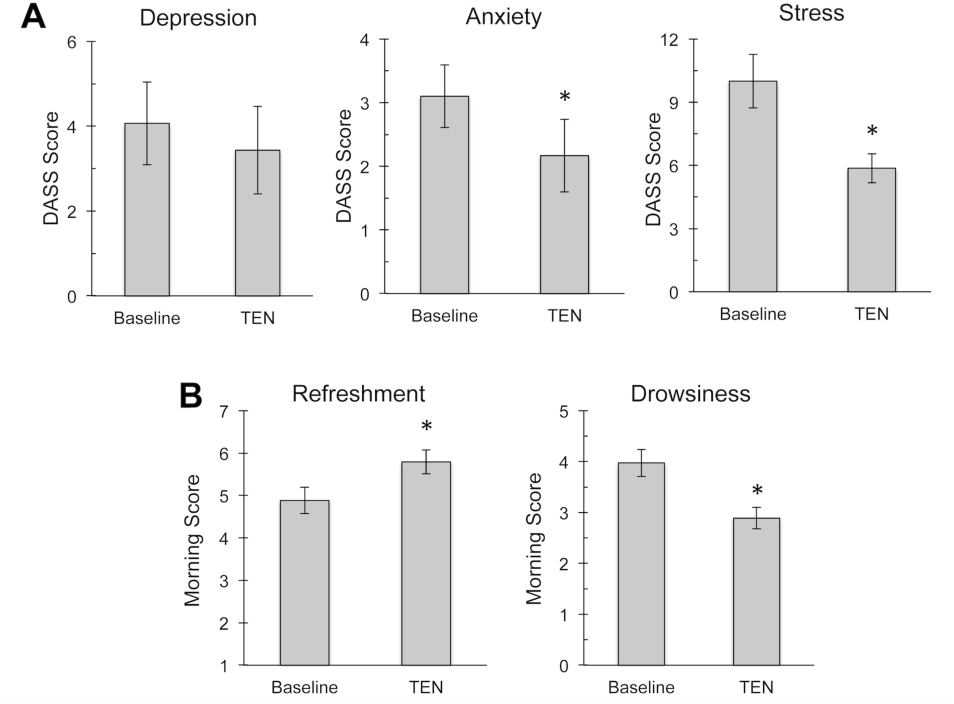
Acute transdermal neuromodulation of trigeminal and cervical afferents prior to sleep onset significantly reduces stress and anxiety. Data obtained from Experiment 1 are shown as a series of histograms. **A**, The histograms illustrate mean scores obtained from the Depression, Anxiety, and Stress Scale (DASS) and show a significant reduction in stress and anxiety following one week of nightly self-administered TEN compared to the baseline week. **B**, The histograms illustrate mean scores obtained on the daily-administered Awakening Drowsiness and Refreshment Scale (see Methods). These data show that nightly TEN significantly increased the feelings associated with refreshment and significantly decreased feelings associated with drowsiness in the mornings compared to baseline. An asterisk indicates a p < 0.05

### Repeated nightly TEN significantly modulates autonomic activity, increases sleep quality, and improves indicators of mental health

Based on our observations in Experiment 1 and our previously described observations [40], we hypothesized that significantly elevated morning mood and rejuvenation is the result of improved sleep quality via TEN dampening sympathetic nervous system and LC activity prior to bedtime (Table 1 and **Figure 2**). Therefore in Experiment 2 we implemented a mixed design (**Figure 1A**) in a different group of participants to determine the repeatability of TEN effects on mood observed in Experiment 1, as well as to determine the impact of TEN treatment (20 min nightly for 7 nights) on sleep quality and sleep patterns, which are known to be regulated by the autonomic nervous system, LC and alterations in norepinephrine.

In Experiment 2, participants completed one week of baseline followed by one week of sham (n = 17) or TEN (n = 18) treatments. Each morning, participants completed the PANAS, rated waking drowsiness and refreshment, reported number of awakenings, and rated overall sleep quality, and at the end of each week, they completed the DASS. Throughout the experiment, to capture objective sleep metrics, participants wore a Philips Actiwatch2, a clinically validated actigraph that reliably tracks sleep wake/cycles (see Methods). Participants also wore a Polar H7 heart rate monitor to sleep each night to record electrocardiogram data (ECG), from which heart rate (HR) and heart rate variability (HRV) metrics were derived.

Building on the findings from Experiment 1, relative to baseline, TEN treatment improved positive affect by 10.86% (t(17) = 4.859, p < 0.001), but failed to alter negative affect (t(17) = −0.237, p = 0.815; Table 2). Further, during the TEN treatment period, participants felt 18.95% less drowsy (t(17) = −4.859, p < 0.001) and 13.33% more refreshed upon waking (t(17) = 4.908, p < 0.001; Table 2). During the sham-treatment period, there were no changes from baseline in affectivity (PANAS) or waking arousal (p > 0.200; Table 2). To examine the impact of TEN on stress, anxiety, and depression symptoms we conducted a series of one-way analyses of variance (ANOVA) comparing DASS score changes from baseline for the TEN‐ and sham-treatment groups. The TEN-treatment group reported a 59.53% greater reduction in stress (F(1, 25) = 4.907 p = 0.036) and 30.68% greater reduction in anxiety (F(1, 25) = 4.392, p = 0.047) compared to the sham-treatment group (**Figure 3A**). The change in depression symptoms were similar across the two groups (F(1, 25) = 0.091, p = 0.766). These results suggest that relative to baseline, TEN-treatment prior to bed improves mood and symptoms of mental health whereas sham treatments do not.

**Table 2.** Summary of results from Experiment 2 illustrating impact of nightly TEN and sham treatment on morning arousal, affect, and sleep quality compared to baseline nights.

|  | Sham Treatment Group |  | TEN Treatment Group |  |
| --- | --- | --- | --- | --- |
|  | Baseline<br>M (SD) | Sham<br>M (SD) | Baseline<br>M (SD) | TEN<br>M (SD) |
| Drowsiness | 4.35 (1.30) | 3.91 (1.15) | 4.69 (1.59) | 3.80 (1.63)* |
| Refreshment | 3.77 (1.06) | 3.89 (0.87) | 3.89 (0.76) | 4.41 (0.78)* |
| Negative Affect | 2.17 (0.58) | 2.30 (0.72) | 2.04 (0.52) | 2.26 (0.64) |
| Positive Affect | 1.40 (0.40) | 1.41 (0.51) | 1.28 (0.28) | 1.27 (.034)* |
| Sleep Quality | 4.58 (0.70) | 4.36 (0.96) | 4.51 (0.90) | 5.01 (0.89)* |
| # of Awakenings | 1.41 (1.23) | 1.20 (1.40) | 1.19 (0.84) | 0.75 (0.62)* |

**FIGURE 3.**
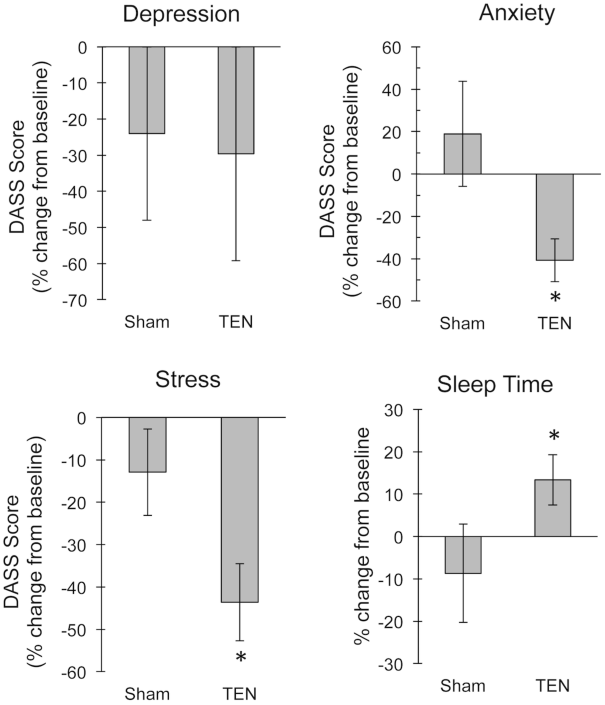
Self-administered trigeminal and cervical afferent neuromodulation prior to sleep onset significantly improves mood and increases sleep time. Data from Experiment 2 are shown as histograms of the percent change in depression, stress, and anxiety as indicated by the DASS, as well as actigraphy-recorded sleep time for TEN and active sham treatment groups compared to baseline. An asterisk indicates a p < 0.05.

Beyond improvements in mood, we observed that TEN produced favorable changes in sleep quality and sleep patterns. Compared to baseline, TEN treatment resulted in a 36.68% reduction in self-reported middle of the night awakenings (t(17) = −3.531, p = 0.003) and an improvement in sleep quality (t(17) = 2.155, p = 0.046; Table 2). There were no differences in middle of night awakenings and sleep quality between the baseline period and sham-treatment period (p > 0.145). Self-reported sleep improvements mirrored sleep cycle changes captured by actigraphy, such that compared to baseline, the TEN-treatment period resulted in significantly increased sleep time (t(14) = 2.255, p = 0.041; **Figure 3B**) and significantly decreased percent time awake (t(14) = −2.329, p = 0.035).

There were no significant differences in sleep time or percent awake time recorded by actigraphy between the sham-treatment period and baseline (all p-values > 0.466). Sleep changes due to TEN uses were also reflected by HRV outcomes. Compared to the baseline period, during the TEN-treatment period participants demonstrated a 4.04% decrease in the relative power of the very low frequency (pVLF) HRV band (t(14) = −2.469, p = 0.027; **Figure 4**) and 9.04% increase in the relative power of the high frequency (pHF) HRV band (t(14) = 2.160, p = 0.049), whereas no changes in other HRV metrics differed between sham treatments and baseline (all p-values > 0.135).

**FIGURE 4.**
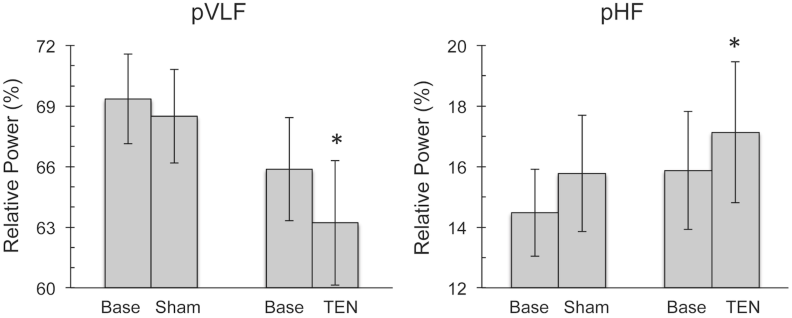
Nightly TEN influences autonomic nervous system activity overnight indicated by changes heart rate variability. Data from Experiment 2 are shown as histograms illustrating the relative powers of the very low-frequency (pVLF) and high-frequency (pHF) bands of heart rate variability (HRV) spectra recorded overnight for one week per treatment condition. The histograms illustrate TEN‐ and active sham-treatment compared to baseline. An asterisk indicates a p < 0.05.

### TEN differentially impacts sleep patterns in a manner dependent on neuromodulation waveform pulse frequency

In Experiment 3 we implemented a randomized double-blinded crossover design (**Figure 1B**) in a different set of participants (n = 25) to determine if a frequency-shift in the neuromodulation waveform parameters used would produce the same impact on sleep quality and mood as we observed in Experiments 1 and 2. We specifically compared the TEN waveform (3 – 11 kHz pulse frequency, 5-7 mA average current amplitude) used in Experiments 1 and 2 to an active sham treatment (n = 10) and a lower frequency TEN treatment (TEN_LF_; 0.5 – 0.75 kHz, < 5 mA average current amplitude; n = 12). In pilot experiments we found TEN_LF_, compared to higher frequency TEN, produced more robust effects on subjective reports of relaxation and physiological indicators of stress suggesting a more pronounced dampening of sympathetic nervous system activity (**Figure S1**). We therefore hypothesized that TEN_LF_ might produce stronger effects on sleep. TEN treatments were grouped as TEN and Control treatments, such that TEN and active sham served as Control treatment groups while TEN_LF_ and TEN comprised the TEN treatment group as specified. Each treatment period lasted one week beginning on consecutive Fridays, with assessments at similar times for each week for each participant. At each of three office visits participants completed the Pittsburg Sleep Quality Index (PSQI) and the DASS. Each morning during the treatment weeks participants completed the Karolinska Sleep Diary (KSD), a validated measure used to capture information about prior night's sleep (see Methods). As in Experiment 2, participants wore the Philips Actiwatch2 throughout the testing period and a Polar H7 heart rate monitor throughout each night.

We first found that TEN produced the same effects on sleep quality and mood compared to control as we observed in Experiments 1 and 2 (data not shown). We next examined the changes that TEN produced on stress, anxiety and depression (DASS) compared to control. We observed that TEN produced a 36.13% greater reduction in anxiety than sham Control (t(20) = −3.406, p = 0.003). There were no perceptible changes in symptoms of stress (t(20) = 0.670, p = 0.510) or depression (t(20) = −1.418, p = 0.172) between the two conditions. Further, across the two conditions (TEN vs active sham Control), there were no changes in self-reported sleep patterns, including sleep quality, time till sleep onset, number of awakenings, and sleep duration. However, the objective actigraph measurements captured changes in sleep/wake cycles such that TEN treatment resulted in 12.12% less wake time (t(19) = −2.381, p = 0.028), 9.50% fewer middle of the night wake-ups (t(19) = −3.781, p = 0.001), and a 9.13% reduction in the percent of wake time (t(19) = −2.241, p = 0.037) compared to sham Control (**Figure 5A**). Despite stability of self-reported sleep pattern data between TEN and control treatments, the data obtained using actigraphy reliably suggest clear improvements in the sleep/wake cycle when using TEN compared to sham Control. We also observed effects reflecting changes in autonomic balance as indicated by significant differences in HR and HRV between TEN and sham control treatment. Relative to active sham Control, TEN decreased the R-R (t(14) = −2.372, p = 0.033), marginally increased HR (t(14) = 1.927, p = 0.075), decreased RMSSD (t(14) = −4.144, p = 0.050), marginally increased the relative power of the LF band of the HRV spectrum (t(14) = 1.878, p = 0.081), and significantly increased the peak LF (t(14) = 3.873, p = 0.002; **Figure 5B**). All other HRV metrics remained stable across the two conditions (all p-values > 0.289).

**FIGURE 5.**
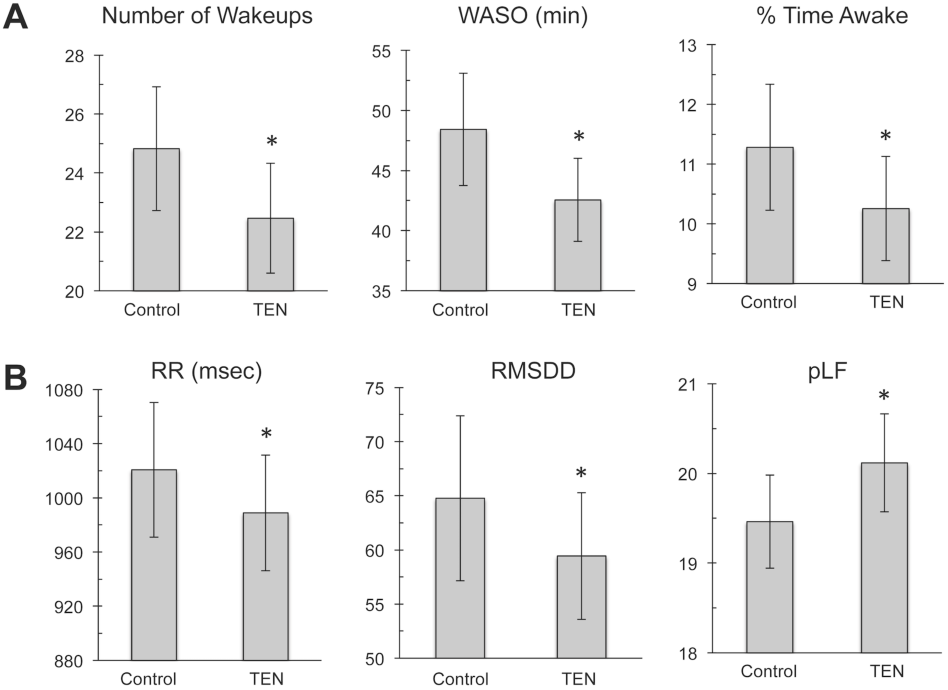
Actigraphy shows that TEN significantly improves sleep quality. Results from Experiment 3 showing differences in actigraphy metrics and HRV are shown for TEN and Control groups as histograms. **A**, The mean number of nightly wakeups, time awake after sleep onset (WASO), and percent time awake recorded using actigraphy illustrate that TEN exerted a significant improvement in sleep quality compared to active sham Controls. **B**, The mean R-R interval and root mean square of the standard deviation (RMSDD) for HRV, as well as the relative power of the low-frequency (pLF) band of HRV spectra are shown for TEN and active sham Control treatments. An asterisk indicates a p < 0.05.

In the subset of participants that received the TEN_LF_ and TEN treatments, we found that TEN_LF_ improved sleep quality and mood outcomes beyond the effects evidenced by TEN in Experiments 1 and 2. Compared to TEN, TEN_LF_ reduced self-reported actiwatch-recorded number of wake-ups (t(9) = 2.796, p = 0.021), wake time after sleep onset (t(9) = 2.808, p = 0.020), and percent of time awake after sleep onset (t(9) = 3.006, p = 0.015; **Figure 6A**). In addition, we observed that TEN_LF_ produced a 32.17% greater reduction in anxiety from baseline than TEN (t(8) = −2.382, p = 0.044), but there were no changes in symptoms of stress (t(8) = 0.928, p = 0.381) or depression (t(8) = − 0.769, p = 0.467; Figure 6B).

**FIGURE 6.**
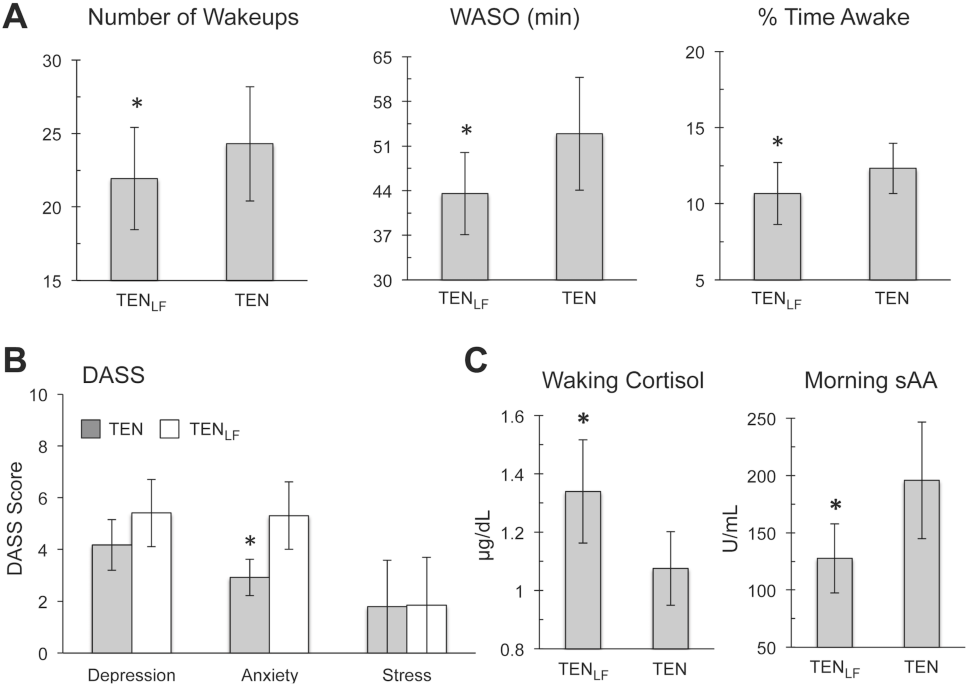
Low-frequency TEN improves the efficacy of trigeminal and cervical afferent modulation for improving sleep quality. Results from Experiment 3 showing differences in actigraphy metrics and HRV are shown for TEN low-frequency (TEN_LF_) and the Control TEN groups as histograms. **A**, The mean number of nightly wakeups, time awake after sleep onset (WASO), and percent time awake recorded using actigraphy illustrate that TEN_LF_ exerted a significant improvement in sleep quality compared to TEN Controls. **B**, Histograms illustrating data obtained from the DASS across treatment groups are illustrated. **C**, The histograms illustrate the results obtained from biochemical quantification of waking cortisol concentrations and α-amylase activity in saliva across treatment conditions. An asterisk indicates a p < 0.05.

### TEN impact on stress and sleep biomarkers, cortisol and salivary α-amylase

The protein enzyme α-amylase is widely recognized as a biochemical marker of sympathetic nervous system activity and sympathoadrenal medullary (SAM) axis activation [42-45]. More specifically, salivary levels of α-amylase directly correlate with plasma norepinephrine (NE) concentrations following the induction of acute stress [42-48]. It has been shown in animals ranging from flies to humans that α-amylase is a reporter of sleep drive as it increases with accumulating sleep debt [49]. To assess the impact of TEN on SAM axis activity as it relates to sleep/wake cycles and daily stress, we examined fluctuations in salivary α-amylase (sAA) and cortisol levels upon waking, in the evening and before bed the final 3 days in each week of the TEN vs TEN_LF_ treatments during Experiment 3 (N = 8, **Figure 1B**). We found that the mean waking levels of sAA were significantly lower in the TEN_LF_ group (TEN_LF_ = 133.78 ± 85.83 U/mL) compared to the TEN group (TEN = 217.33 ± 134.23 U/mL); t(5) = −3.524, p = 0.017; **Figure 6C**). There were no differences between TEN treatments in afternoon and bedtime sAA (p > 0.5). These data indicate that compared to TEN, TEN_LF_ produced changes reflecting lower levels of the sleep debt and stress upon awakening.

While sAA reflects NE levels and acute SAM axis activity, cortisol is another biomarker, which is under complex hormonal control of the hypothalamic-pituitary-adrenal (HPA) axis. Elevated salivary cortisol levels prior to bedtime are a predictor of insomnia often related to hyperarousal, metabolic dysfunction and other mood disorders, but its causal importance in poor sleep remains unclear [50-53]. Reflective of adrenal fatigue however, insomnia and poor sleep have been shown to significantly dampen the cortisol awakening response (CAR), which is a specific phase of the diurnal cortisol rhythm [54, 55]. We observed no differences in afternoon or bedtime cortisol levels across the treatment groups (all p-values > 0.5). We did find that when participants used TEN_LF_ prior to bed they exhibited a stronger CAR, a significant 26.3% higher salivary cortisol concentration compared to when they used TEN (TEN_LF_ waking cortisol concentration = 1.44 ± 0.41 μg/dL, TEN waking cortisol concentration = 1.14 ± 0.322 μg/dL; t(5) = 2.850, p = 0.036; **Figure 6C**). These particular observations indicate that TEN_LF_ produced more robust effects than TEN on the restorative features of sleep by significantly improving the quality of the sleep/wake cycle.

## DISCUSSION

The ascending reticular activating system (RAS) represents a collection of brain stem nuclei and neuromodulatory projections, such as noradrenergic radiations from the locus coeruleus (LC) and cholinergic axons from the pedunculopontine nucleus (PPN) to higher brain circuits including the thalamus, hippocampus, amygdala, and vast regions of cortex. These ascending networks are known to provide strong neurophysiological control of consciousness, sleep/wake cycles, attention, alertness, other aspects of cognition including behavioral flexibility [1, 15, 28, 56-60]. Starting from the periphery, primary trigeminal and cervical afferent inputs to the trigeminal sensory nuclear complex (TSNC) from sensory and proprioceptive fibers located on the sides and front of the face, oral cavity, and anterior and posterior cervical regions of the neck provide robust disynaptic and polysynaptic regulation of LC and PPN neuron activity. Demonstrating functional effects on ascending RAS circuitry, we have shown that high-frequency (7 – 11 kHz) transdermal electrical neuromodulation (TEN) of trigeminal and cervical afferents can rapidly suppress psycho‐ and neuro-physiological arousal as indicated by a significant dampening of sympathetic nervous system activity and suppression of noradrenergic and sympathomedulary (SAM) axis biomarkers in response to an acute stress challenge [40]. Given this context, in the present report we tested the hypothesis that self-administered, nightly TEN prior to bedtime would improve sleep quality and mood. Our conclusions based on data and observations gathered from the three independent experiments described in this study are that nightly TEN does cause a significant improvement in the quality of sleep and mood across weeklong timescales. Below we discuss more specific details related to these findings and their implications for optimizing mental health and human performance.

### Interfacing with the trigeminal sensory nuclear complex to functionally regulate ascending neuromodulatory systems

There is a growing need for neural interfaces capable of modulating brain function and behavior in the least invasive manner possible. For the past several years we have explored the potential of utilizing peripheral nerve pathways to tap into brain function. Specifically, we have been focused on developing novel approaches to trigeminal and cervical nerve modulation (**Figure 7**). Each of these nerves, as well as other peripheral and cranial nerves offer unique (and overlapping) structural inroads to deep-brain nuclei responsible for regulating psychophysiological arousal, attention, neurobehavioral responses, and physiological performance through ascending and descending circuitry. Based on the evidence we have accumulated from observations in thousands of participants we have strong beliefs that modulation of trigeminal and cervical nerve pathways can offer a chemical-free way of regulating autonomic arousal to improve and optimize mental health and human performance. With continued research and development, there will be some near-term opportunities to offer treatments for some of the most broadly debilitating mental health disorders facing the modern world using transdermal trigeminal and cervical nerve modulation. Attention must be paid to what behavioral paradigms and treatment regimes best support implementation for a specific indication, but the core neurobiological outcomes strongly suggest that treatment of daily stress, anxiety, and comorbid sleep disturbances would be efficient and practical using TEN of the trigeminal and cervical nerves. Likewise, we believe that there will be an opportunity to modulate certain cognitive processes using the same or similar bottom-up modulation approaches via cranial and cervical nerves to induce higher-order plasticity in cortical and subcortical brain regions. For example, pairing of bottom-up NE and ACh modulation using TEN of trigeminal and cervical nerves with top-down cortical processing that occurs during skill training could prove to be a powerful method of accelerating learning or acquiring expert proficiency. Future studies using peripheral nerve modulation to enhance plasticity will directly address this possibility.

**FIGURE 7.**
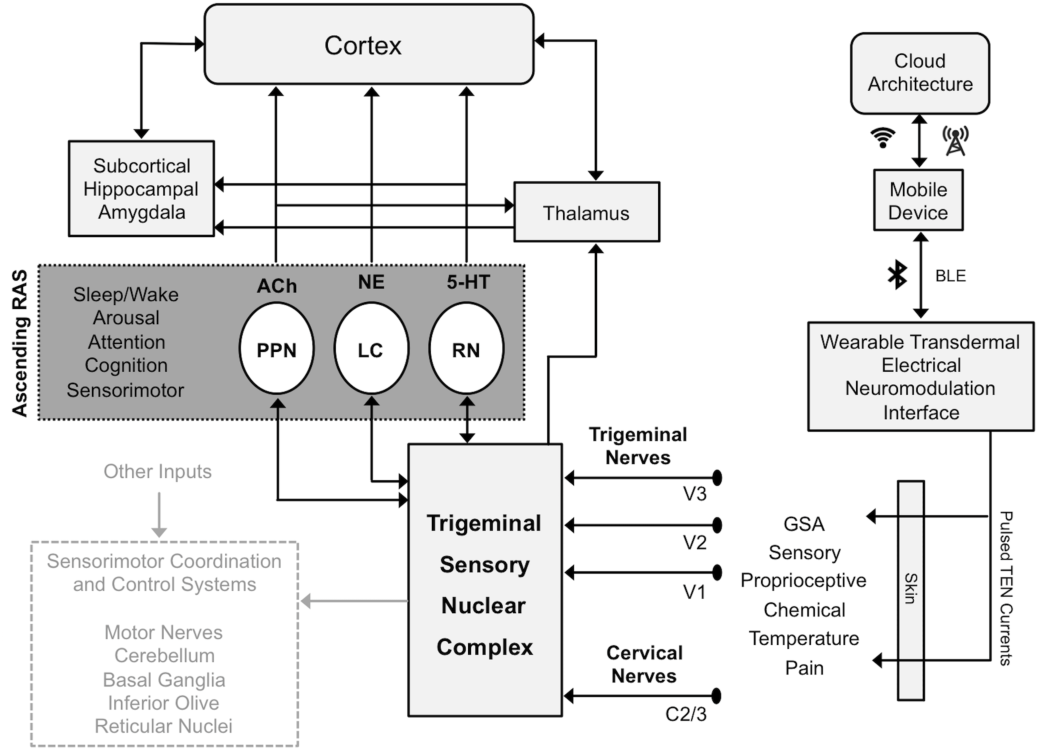
Interfacing with the ascending reticular activating system through the trigeminal sensory nuclear complex. The illustration depicts the approach to modulating trigeminal and cervical nerve afferents, which provide primary sensory inputs to the trigeminal sensory nuclear complex (TSNC). Similar to previously described methods [40] volunteers modulated their trigeminal and cervical nerves using a wearable transdermal neurostimulator for 20 min each night prior to sleep onset. The TSNC coordinates motor behaviors by sending projections through a series of ascending and descending pathways communicating with various targets (greyed out). An emphasis is placed however on the ascending reticular activating system (RAS), which includes the pedunculopontine nucleus (PPN), the locus coeruleus (LC), and raphe nuclei (RN). These pathways transmit neuromodulatory acetylcholine (ACh), norepinephrine (NE), and serotonin (5-HT) signals to higher-order brain structures to gate attention and regulate awareness, arousal, and sleep. As discussed in the text, the TSNC and its inputs have been shown provide functional connections to the LC and PPN and thereby provides a path for the ability of trigeminal and cervical nerve modulation to influence sleep as described in the present study. These pathways also open the possibility of regulating attention, cognition, and sensorimotor behaviors using trigeminal and cervical nerve modulation paired with training or other approaches.

The use of TEN to modulate neural activity is supported by a long history of safety obtained over four decades. There are numerous methods and devices intended for modulating peripheral nerve structures using transcutaneous delivery of voltage/current waveforms from electrodes applied to various locations on the body. These devices such as transcutaneous electrical nerve stimulation (TENS), powered muscle stimulation (PMS), electrical muscle stimulation (EMS) and others have amassed such a high degree of physical safety that they have been moved to an over-the-counter product rather than a medical device requiring a prescription depending on the intended use and design characteristics. For example, legally marketed electrical nerve stimulation devices are already commercially available and have output levels far greater than the ones we implemented here. These devices intended for over-the-counter cosmetic applications of TENS target similar anatomical regions and nerve targets such as the trigeminal nerve. One example is an over-the-counter cosmetic TENS device (Bio-medical Research Face), which is designed to target the trigeminal nerve and provide neuromuscular electrical stimulation (NMES) to encourage facial rejuvenation for aesthetic purposes. A recent study examined the safety and efficacy of this device at a peak current intensity (35 mA) that was nearly twice the one used in our study when used five days per week for 20 minutes each day for 12 weeks [61]. There were no significant adverse events in this study and the only reported side effects were minor skin redness following stimulation, which disappeared with 10-20 minutes following use [61]. Another device, which modulates supraorbital branches of the trigeminal nerve to treat headache has also demonstrated a high safety threshold when used daily for multiple weeks [37]. Other reports using trigeminal nerve stimulation for the treatment of epilepsy, depression, and other disorders have likewise shown a high degree of safety [34-36, 62-64]. Although there is a high degree of confidence in the safety of trigeminal nerve modulation, caution is always warranted when delivering electrical currents to the human body and we advise investigators to learn and implement safe practices using qualified devices.

### The effects of nightly self-administered TEN on mood and sleep quality in a natural environment

We examined the effects of trigeminal and cervical TEN on sleep quality and mood in healthy volunteers having mild to moderate sleep disturbances (PSQI > 5). Various tools and sensors were deployed in real world environments such that data could be acquired from within the homes of volunteers under baseline conditions or following self-administered TEN or sham treatments in a double-blinded fashion depending on the experimental protocol (see Methods). Through a cloud computing architecture, we were able to remotely monitor treatment sessions in order to ensure compliance throughout the study (**Figure 7**). We found that one-week of nightly self-administered TEN of trigeminal and cervical afferents (20 min/night) significantly improved mood and sleep quality in home environments. In Experiment 1, we found that one-week of nightly TEN (20 min/night) significantly reduced stress and anxiety compared to baseline DASS scores (**Figure 2**). This improvement in mental health indicators was accompanied by a significant reduction in morning drowsiness and significant increase in morning refreshment following TEN of trigeminal and cervical afferents compared to baseline mornings (**Table 1** and **Figure 2**). TEN also caused a significant increase in positive affect and a significant decrease in negative affect according to the PANAS administered on mornings after real treatments compared to sham treatments or baseline nights (**Tables 1** and **2**). In Experiments 2 and 3 we more closely inspected how TEN affected sleep quality using actigraphy. We found that compared to active sham treatment, TEN significantly increased the amount of sleep time and significantly reduced the time awake after sleep onset (WASO; **Figures 3** and **5**). As discussed in more detail below, a low-frequency (0.5 – 0.75 kHz) shifted TEN waveform (TEN_LF_) caused a significant improvement in sleep quality (**Figure 6**) compared to the standard high-frequency TEN waveform we used that has been previously described (see *Transdermal neuromodulation waveform considerations* below) [40]. We suspect that the reduction in stress and anxiety, as well as the improved positive affect on weeklong timescales was a secondary outcome of the primary effect that TEN of trigeminal and cervical nerves had on improving sleep quality in a natural or home environment. Future studies examining how TEN of trigeminal and cervical nerves affects sleep structure using polysomnography will inform the design of enhanced approaches to improving general sleep quality, as well as to optimizing the time spent asleep or resting. Such approaches will have a significant impact on our ability to improve daily mental health and restore or enhance human performance.

### The effects of TEN on autonomic activity during sleep

There is known to be a tight coupling of central nervous system sleep regulation and the activity of the autonomic nervous system where heart-rate control under different stages of sleep in healthy subjects is apparent [65]. During REM sleep heart rate is known to increase [66]. The relative power of the VLF and HF components of the HRV spectra have been shown to be dominant during REM sleep and deep sleep respectively [66, 67]. There is a marked decrease in the relative power of the VLF and LF HRV spectral bands during deep sleep and an increase during REM sleep compared to HF [66, 67]. Since we captured data related to overall sleep quality rather than sleep structure, we cannot draw strong conclusions related to the effects of TEN on HRV as it relates to the stages of sleep. We do conclude however that given the specific changes observed in the HRV power spectra, compared to controls both TEN and TEN_LF_ produce significant alterations to sleep/wake cycling throughout the night. This is somewhat expected given anatomical connectivity and the functional ability of trigeminal nerve and TSNC activity to modulate the activity of the PPN and LC, which are both known to be central regulators of sleep patterns including REM sleep and waking behaviors [1, 27-30]. More detailed investigations examining sleep structure with additional polysomnography measures will be required to provide further insight into how TEN affects the specific stages of sleep.

### Transdermal neuromodulation waveform considerations

For reasons discussed in greater detail elsewhere [40], we originally developed TEN methods utilizing pulse frequencies in the 2 – 20 kHz range. Stimulating neuronal activity in the LC and PPN at specific frequencies can trigger state changes in sleep/wake cycles [1, 12-14]. Therefore, we questioned whether altering the pulse parameters of high-frequency TEN waveforms would produce different effects on sleep quality or mood. We specifically chose to lower the frequency of the TEN waveforms by about an order of magnitude such that we maintained a neuromodulation frequency above 0.12 kHz. We did this because it had previously been shown that transcutaneous modulation of supraorbital branches of trigeminal nerve afferents at 0.12 kHz induces sedative-like states in humans whereas 0.025 kHz did not [39]. This observation indicated to us that frequencies higher than 0.12 kHz could be even more effective at inducing deep states of relaxation and perhaps, alter sleep quality. Based therefore on our previous observations [40] and those made by Piquet and colleagues (2011) we investigated the effects of low-frequency TEN (TEN_LF_; 0.5 – 0.75 kHz) waveforms that utilized a base neuromodulation frequency < 7 kHz and > 0.12 kHz on sleep quality and mood to compare and contrast them to those obtained with high-frequency TEN waveforms. We found that TEN_LF_ caused a significant reduction in the number of nightly wake-ups, a significant reduction in WASO, and a significant reduction in the percentage of time spent awake during the night compared to high-frequency TEN, which itself was significantly effective at improving sleep quality compared to sham and baseline observations (**Figures 3**, **5**, and **6**). Interestingly, the effects of TEN_LF_ on sleep were observed at lower average current amplitudes than those used for standard TEN treatments (see Methods). We also found that TEN_LF_ produced a significant reduction in DASS anxiety scores compared to high-frequency TEN (**Figure 6**). There was additional biochemical evidence that TEN_LF_ was significantly better at improving restorative sleep than high-frequency TEN.

The protein enzyme α-amylase is as a biochemical marker of noradrenergic activity and sympathoadrenal medullary (SAM) axis activation [42-45]. Further, it has been shown that α-amylase is a reporter of sleep drive as it increases with accumulating sleep debt [49]. Compared to high-frequency TEN treatments, the waking levels of salivary α-amylase (sAA) were significantly lower following treatment with TEN_LF_ waveforms prior to bed (**Figure 6C**). Cortisol is another stress biomarker, which is under the complex control of the hypothalamic-pituitary-adrenal (HPA) axis. Insomnia and sleep debt can lead to adrenal fatigue and significantly dampen the cortisol awakening response (CAR), which specific phase of the diurnal cortisol rhythm [54, 55]. We did find that when participants used TEN_LF_ prior to bed they exhibited a significantly stronger CAR compared to when they used TEN (**Figure 6C**). These particular observations indicate that TEN_LF_ produced more robust effects than TEN on the restorative features of sleep by significantly improving the quality of the sleep/wake cycle, as well as altering morning levels of stress biomarkers and sleep debt. In other words, TEN produces a positive impact on mood and sleep quality in a frequency-dependent manner. These observations illustrate the need for systematic research exploring how discrete waveform parameters, such as pulse frequency affect physiological and biochemical responses in humans. Studies in humans can be supported by studies in other animal models where understanding circuit level input-output relationships by conducting electrophysiological recordings from different brain regions is more feasible.

### Potential implications for the field noninvasive brain stimulation

Projections from the TSNC to brain structures like the LC and PPN provide a foundation for the formulation of alternative hypotheses that oppose conventional views on top-down mechanisms of action underlying “transcranial” electrical stimulation (tES) approaches. We believe that noninvasive RAS modulation via the TSNC and other ascending cranial/cervical neural (sensory) pathways offer a solid framework for explaining many of the observations made using the passage of low-intensity direct currents (≤ 5 mA peak) across the skin and presumably the scalp, dura, and cerebrospinal fluid before influencing the cortical surface (for example, as has been proposed to be a mechanism for transcranial direct current stimulation or tDCS). We cannot imagine a situation where the ascending endogenous neuromodulatory pathways carrying sensory information via cranial nerves do not contribute a significant degree to the observations made using tDCS and some other tES methods. Our studies as well as the studies of many others [16, 18, 23, 32, 33, 39, 62, 63, 68-77] on electrical neuromodulation or stimulation of cranial nerves indicate that investigators applying electrical currents (DC, AC, or other) to the skin of the head should consider the impact of bottom-up pathways as illustrated in **Figure 7**. The possible involvement of bottom-up pathways has been almost entirely overlooked by the scientific community that implements tES. Modulation of afferent pathways via cranial nerves, cervical spinal nerves, and other peripheral systems can explain many if not most of the outcomes observed in studies implementing tES approaches. A simple inspection of neuroanatomy illustrates that cranial nerves carrying afferent signals to the brain stem and RAS structures cannot be avoided when placing electrodes on the head. Further, electrical stimulation of the dura has been shown to activate neurons in the TSNC [77], which then project to numerous ascending neuromodulatory nuclei.

Highlighting mechanistic inconsistencies regarding tES approaches, explanations underlying the top-down modulation of PFC function using tDCS have led to a high-degree of variability and uncertainty regarding the robustness of behavioral, neurophysiological, and cognitive outcomes [78-80]. Modulation of the LC activity and noradrenergic signaling, which can be achieved via cranial nerve modulation as previously discussed [40], has been shown to effect: vigilance and sustained attention [57, 81], alertness and arousal [10, 82, 83], working memory and decision making [15, 84, 85], the amplitude of motor evoked potentials [86, 87], brain oscillations [88-90], sensory gating and processing [10, 91, 92], and others. These same outcomes have also been claimed via direct modulation of cortical circuits by numerous tES investigations.

Due to spatial and temporal resolution limits it seems that fMRI, EEG, and other neurophysiological or psychological measurements of slowly (hundreds of milliseconds to seconds) evolving biological processes would have difficulty resolving direct cortical effects versus those delayed by transmission at only a couple synapses, which happen to mediate some of the most powerful endogenous neuromodulatory actions upon the human brain. For example, transcranial alternating current stimulation (tACS) of the prefrontal cortex was recently shown to exert an effect on dreaming. It was reported that 25 - 40 Hz tACS delivered to the prefrontal cortex triggered lucid dreaming and increased conscious self-awareness during dreaming [93]. However trigeminal nerve entrainment of PPN neurons, which as previously mentioned are known to regulate REM sleep at those frequencies [1, 12], could also explain the effects of tACS on dream (sleep) and conscious self-awareness. Thus to challenge the dogma, we urge the neuromodulation community to explore cranial nerve modulation as an alternative hypothesis capable of mechanistically explaining many of the outcomes observed in response to conventional tES.

### Conclusion

For the past decade evidence has been steadily accumulating that the trigeminal nerves provide a unique opportunity to tap deep into human brain function. The embedded connectivity of the TSNC with RAS structures like the LC and PPN make trigeminal nerves a high priority peripheral nerve target when using neuromodulation to regulate stress, attention, arousal, and sleep/wake cycles (Figure 7). We previously reported TEN of trigeminal and cervical nerves can significantly dampen sympathetic nervous system activity in response to acute stress [40]. In the present study we showed that repeated, nightly TEN significantly improves sleep quality and significantly reduces stress and anxiety over the course of a week in healthy volunteers. Others have shown that transcutaneous trigeminal neurostimulation is effective at treating drugresistant epilepsy and depression [31, 35, 36]. There is also preliminary clinical evidence that trigeminal neurostimulation has therapeutic value in managing the symptoms of PTSD [32], generalized anxiety disorder [33], and ADHD [34], Presently there is a surging interest in harnessing the capability of peripheral nerves to induce brain plasticity. Ongoing and forthcoming studies will lead to new knowledge and a refinement in our ability to regulate brain function and behavior through peripheral nerve modulation. We conclude that trigeminal and cervical nerve modulation of ascending RAS circuits will become increasingly useful for optimizing human performance by regulating states of consciousness, such as sleep/wake cycles, neuro‐ and psychophysiological arousal, and attention or vigilance.

## METHODS

### Participants

All experimental procedures were conducted on human volunteers using protocols approved by an independent Institutional Review Board (Solutions IRB, Little Rock, AR), All subjects provided written informed consent prior to experimentation. Exclusion criteria were as follows: diagnosed sleep disorder, actively medicated for sleep difficulties, neurological or psychiatric disorder, cranial or facial metal plate or screw implants, severe face or head trauma, recent concussion or brain injury, recently hospitalized for surgery/illness, high blood pressure, heart disease, diabetes, pregnant, acute eczema on the scalp, and uncorrectable vision or hearing. In Experiment 3, we examined participants with moderate sleep impairment, screened as presenting with a score of 5 or greater on the Pittsburg Sleep Quality Index [94], A subset of these participants was selected to provide saliva samples for biometric assays for which several additional exclusion criteria applied: nicotine use, recreational drug use, and dental work during the experimental period. Further, to minimize experimental complexity due to health and due to female hormone fluctuations, which impact the biometric markers being assayed, age range was restricted to 20 to 40 years old and males were oversampled.

Experiment 1 was designed to examine the impact of TEN prior to bedtime on acute and long-term mood, assessed each morning with the Positive and Negative Affectivity Scale (PANAS) [95] and ratings of drowsiness and refreshment, and at the end of each week with the Depression, Anxiety and Stress Scale (DASS). Participants completed a weeklong baseline assessment period followed by a weeklong TEN-treatment period. We enrolled 43 participants, 5 of which were withdrawn for failing to comply with study procedures. Twenty-one participants were men, 17 were women, their ages ranged from 20 to 62 (mean age = 29.68 ± 10.88 years), and 67.6% were white, 23.5% were Asian, 2.9% were Black and 5.9% were Hispanic.

Experiment 2 was designed to examine the impact of before bed TEN treatments on sleep patterns and sleep quality, using actigraphy, heart rate monitoring, self-reported number of wake-ups, ratings of sleep quality and the same mood assessment administration as Experiment 1. We enrolled 42 participants, 6 of whom withdrew due to scheduling conflicts and time commitments and 1 that was removed for failing to comply with study procedures. Of these participants, 62.1% were female; their ages ranged from 19 to 59 (mean age = 27.66 ± 9.9) and 62.1% were white, 17.2% were Asian, 17.2% were Black and 3.4% were Hispanic.

Experiment 3 was designed to directly compare the impact of bedtime TEN treatments to Sham treatments (active or inactive) on sleep patterns and sleep quality. Experiment 3 employed the same metrics as Experiment 2, with the addition of administration of the DASS during the initial training visit and the PSQI at each visit. In addition, a subset of male participants (N = 7) were randomly selected to provide waking, afternoon and before bed saliva samples during the 3 final days of each treatment week. The enrolled sample comprised 27 participants, 2 of which were removed for failing to comply with study procedures or failing to report for scheduled appointments. Ninety-three percent of participants were male, their average age was 32 years old (SD ± 5.29) and 79% were Caucasian, 11% were Black, 7% were Asian and 4% were Hispanic. As mentioned above, all participants reported active sleep disturbance with PSQI scores ranging from 5 to 15 (mean = 9.56 ± 2.55) and 50% of the sample had a score of 9 or below. In addition, participants reported trouble falling asleep or trouble staying asleep at least once or twice a week, with more than 50% of participants on average taking more than 30 minutes to fall asleep each night (mean = 34.63 ± 24.02).

### Transdermal electrical neuromodulation

The transdermal electrical neuromodulation (TEN) waveform developed for use in these experiments was a pulse-modulated (3 – 11 kHz), biphasic electrical current producing average amplitudes of 5 – 7 mA for 20 min as similarly described in a previous report [40]. The sham waveform was an active stimulation control, in which the waveform parameters outlined above were active for only the first and last 30 secs of the 20 min treatment, providing a significantly lower dosage of the TEN treatment and offering skin sensations that mimicked real treatment. Participants were not able to distinguish between real TEN and sham waveforms. In a subset of participants in Experiment 3, the real TEN waveform described above was compared to a lower-frequency TEN waveform (TEN_LF_; pulse frequency 0.50 – 0.75 kHz, < 5 mA average amplitude), which in lab tests produced more pronounced changes on autonomic nervous system balance (Figure S1). During both the real TEN and sham stimulus protocols, subjects were instructed to adjust the current output of a wearable TEN device (Thync, Inc., Los Gatos, CA) using an iPod touch connected to the device over a Bluetooth low energy network such that it was comfortable. TEN and sham waveforms were delivered to the right temple (10/20 site F8) and base of the neck (5 cm below the inion) using custom-designed electrodes comprising a hydrogel material and a conductive Ag/AgCl film secured to the wearable TEN device. The anterior electrode positioned over F8 was a 4.9 cm^2^ oval having a major axis of 2.75 cm and a minor axis of 2.25 cm while the posterior electrode was a 12.5 cm^2^ rectangle with a length of 5 cm and a height of 2.5 cm. The average current density was < 2 mA/cm^2^ at all times to keep in accordance with general safety practices to prevent any damage to the skin. Across all experiments, participants were trained to use the TEN device on their first visit and were instructed to use within 30 min prior to bed. In Experiments 2 and 3, subjects were assigned to experimental conditions using a randomization method or a counterbalancing approach, and subjects and researchers were always kept blind to experimental conditions.

### Mood Scales and Sleep Surveys

We utilized several scales and schedules to evaluate the impact of TEN on mood and sleep quality. The self-reported scales used in the present study are described below. The Awakening Drowsiness and Refreshment scale was a 5-item self-report measure that is administered in the mornings, within a half hour of waking, to assess feelings of awakening drowsiness (tired and lethargic) and refreshment (refreshed, alert and clear-headed). Items were rated on a 1 - low to 10 - high scale. Across baseline and TEN treatment periods for Experiments 1 and 2 internal consistencies ranged from 0.72 to 0.83 for drowsiness and 0.72 to 0.89 for refreshed. The Positive Affect and Negative Affect Schedule (PANAS) [95] is a clinically validated scale that comprises two 10-item scales: positive affectivity and negative affectivity. Participants rate "how they feel right now” on a 5 point scale: 1 - very slightly/not at all to 5 - extremely. Across all phases of experiments 1 and 2 internal consistencies ranged from 0.94 to 0.96 for positive affectivity and 0.65 to 0.93 for negative affectivity. The Karolinska Sleep Diary (KSD) [96] is a self-report questionnaire administered in the mornings that captures information about the prior nights sleep (e.g., time in bed, time asleep, number of wake-ups, and ratings of degree of dreaming, calm sleep, and sleep quality). This questionnaire has been validated against polysomnography and is correlated with objective EEG sleep metrics (for example, the amount of slow-wave sleep and sleep efficiency). The Depression, Anxiety, Stress Scale (DASS) [97, 98] is a reliable clinically validated self-report measure that comprises 42 negative emotional symptoms. Each item is rated on a 4-point severity/frequency scale and indexed to the past week. Scores for the Depression, Anxiety and Stress scales are determined by summing the scores for the relevant 14 items. Reliabilities on the subscales ranged from 0.53 to 0.95 across all baseline and treatment weeks for all experiments. The Pittsburgh Sleep Quality Index (PSQI) [94] is a 20-item assessment that gauges overall sleep patterns over the past month, including duration of sleep, type of sleep disturbance, sleep latency, sleep efficiency, overall sleep quality, and day dysfunction due to sleepiness. The PSQI was administered in Experiment 3. The baseline administration was indexed to the past month, and subsequent assessments were indexed to past week, capturing sleep changes due to TEN or sham treatment. Scores for the PSQI were calculated according the algorithm outlined by Buysse and colleagues (1989).

### Actigraphy

In Experiments 2 and 3, to monitor sleep/wake cycles, participants wore the clinically validated Phillips Respironics Actiwatch 2 (Phillips Healthcare) on their non-dominant wrist throughout the course of the study. The Actiwatch 2 was equipped with solid-state piezoelectric accelerometer to capture actigraph data at 32 Hz and event markers were used to capture bedtime and morning waking. The Actiwatch 2 has been shown to reliably tracks sleep/wake cycles and calculates duration, wake after sleep onset (WASO), sleep time, wake time, number of wake-ups, and percent sleep/wake [99, 100].

### Heart rate variability

In Experiments 2 and 3, we acquired cardiac output using the Polar H7 heart rate monitor. Recordings started immediately prior to bed, after sham/TEN use, and were terminated upon waking. During participants first office visit they were trained according to manufacturers instructions on preparation and usage of the device. Electrocardiogram (ECG) data were live streamed to the HRV Logger application, which converted the ECG to R-R data that was exported daily to secure servers. We had previously confirmed that HRV Logger recorded data accurately using independent HRV analyses conducted with Kubios and Matlab. We computed an average nightly HR, R-R interval, standard deviation of the normal-to-normal heartbeat (SDNN), and root mean square of the standard deviation (RMSSD). Nightly R-R data were transformed into frequency domain data to examine the relative power of the very low-frequency (pVLF; 0.0033 – 0.04 Hz), low-frequency (pLF; 0.04–0.15 Hz) and high-frequency (pHF; 0.15–0.4 Hz) bands of the HRV spectra, including peak low frequency (pkLF), peak high frequency (pkHF) and the LF/HF ratio.

### Salivary collection and biomarker assays

In Experiment 3, participants provided awakening, afternoon and bedtime saliva samples Tuesday, Wednesday and Thursday of each week of the study. Participants were instructed not to eat or brush their teeth within an hour of providing a sample. Timing of saliva collection varied across participants, but participants provided samples at the same times each day. All morning samples were taken immediately upon waking since this has been shown to reduce variability in assaying the cortisol awakening response [101]. Afternoon saliva samples were taken between 3 PM and 5 PM, and bedtime samples were taken within 30 minutes before bed, which was just prior to TES usage. Salvia was collected via that passive drool method. As per manufacturer’s instructions (SalivaBio, Inc., State College, PA), saliva is pooled at the front of the mouth and eased through a tube, centered on the lips, directly into a cryovial. Saliva samples were immediately stored at –20 °C and were transported back to the testing facility on ice, from there saliva samples were sent to Salimetrics, LLC (State College, PA) where ELISA methods were employed to assess α-amylase (Salimetrics 1–1902) and cortisol levels (Salimetrics 3002). Salivary α-amylase is a biomarker for sleep in humans and is widely recognized as a biochemical marker of sympathetic nervous system activity and sympathoadrenal medullary (SAM) axis activation. The glucocorticoid hormone cortisol has a diurnal rhythm, which peaks within 45 of waking. The cortisol awakening response (CAR) is sensitive to sleep disturbances where poor sleep correlates with a blunted CAR [54, 55].

### Statistical analyses

Given the quick but steep learning curve with fitting the Thync device and given that during training sessions participants received a TEN treatment, we made an a priori decision to exclude the first baseline and TEN-treatment day from all analyses in Experiments 1 and 2. All statistical analyses were completed with IBM SPSS Statistics Software (IBM Corporation, Armonk, NY). In Experiments 1 and 2, unless otherwise stated, all statistical analyses were conducted using paired sample t-tests. In Experiment 3, order effects were examined using two-way repeated measures analyses of variance (RMANOVA). All tests were not significant and were followed by a series of one-way RMANOVA. Across all studies, metrics that were assessed daily were averaged separately across the each assessment period (baseline/TEN/Sham). Participants with fewer than 3 days of data on any metric for any phase of an experiment were considered to have incomplete data on the given metric. Missing or incomplete data were deleted listwise. Thresholds for statistical significance were set at p < 0.05. All data reported and shown are mean ± SD.

## AUTHOR CONTRIBUTIONS

A.M.B. and W.J.T. designed the experiments. A.M.B., H.M.M., R.S.S., and L.A. conducted the experiments. A.M.B., H.M., R.S., L.A., and W.J.T. analyzed data and assisted in the preparation of figures. A.M.B. and W.J.T. prepared and edited of the manuscript.

## ACKNOWLEDGEMENTS

We thank Drs. Sumon Pal and Jonathan Charlesworth for programming the transdermal electrical neuromodulation waveforms. We are in particular gracious to Dr. Sumon Pal for his insight, development of TEN waveforms, and collaborative assistance in pilot studies. We would also like to thank Dr. Dan Wetmore for his critical feedback and insights throughout the planning and execution of the study. We thank all the other members of Thync, Inc., for their invaluable technical and operational support, which was critical to conducting the study. We also thank Thync, Inc., for providing the funding and resources required to conduct this study.

## SUPPLEMENTAL INFORMATION

**FIGURE S1.**
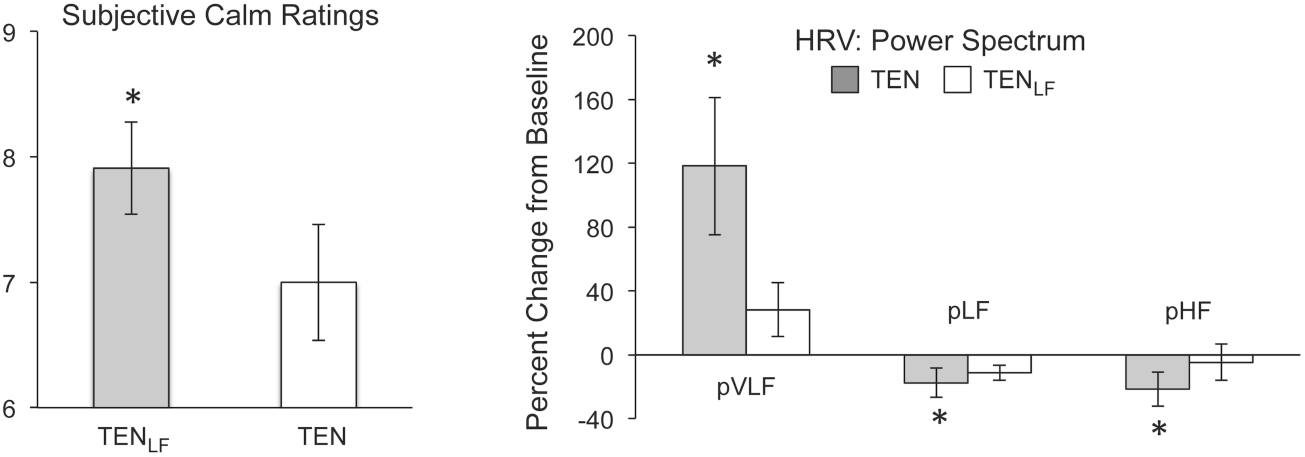
The effects of a low-frequency TEN waveform on subjective relaxation and heart rate variability.

